## Supplementary figures and images for "Spatial Transcriptomics of Meningeal Inflammation Reveals Inflammatory Gene Signatures in Adjacent Brain Parenchyma"

### Supplementary Figure 1

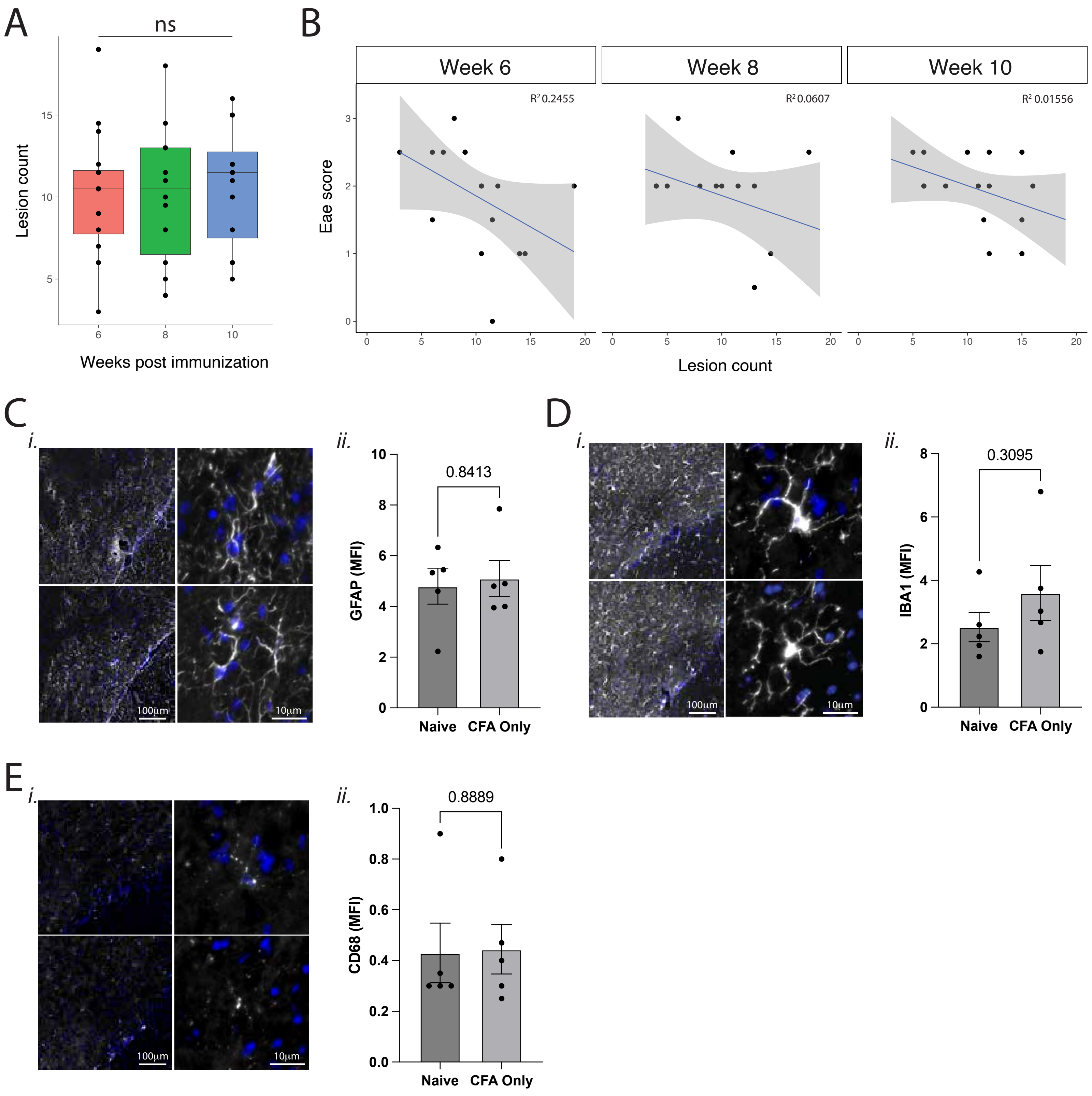

### Supplementary Figure 2

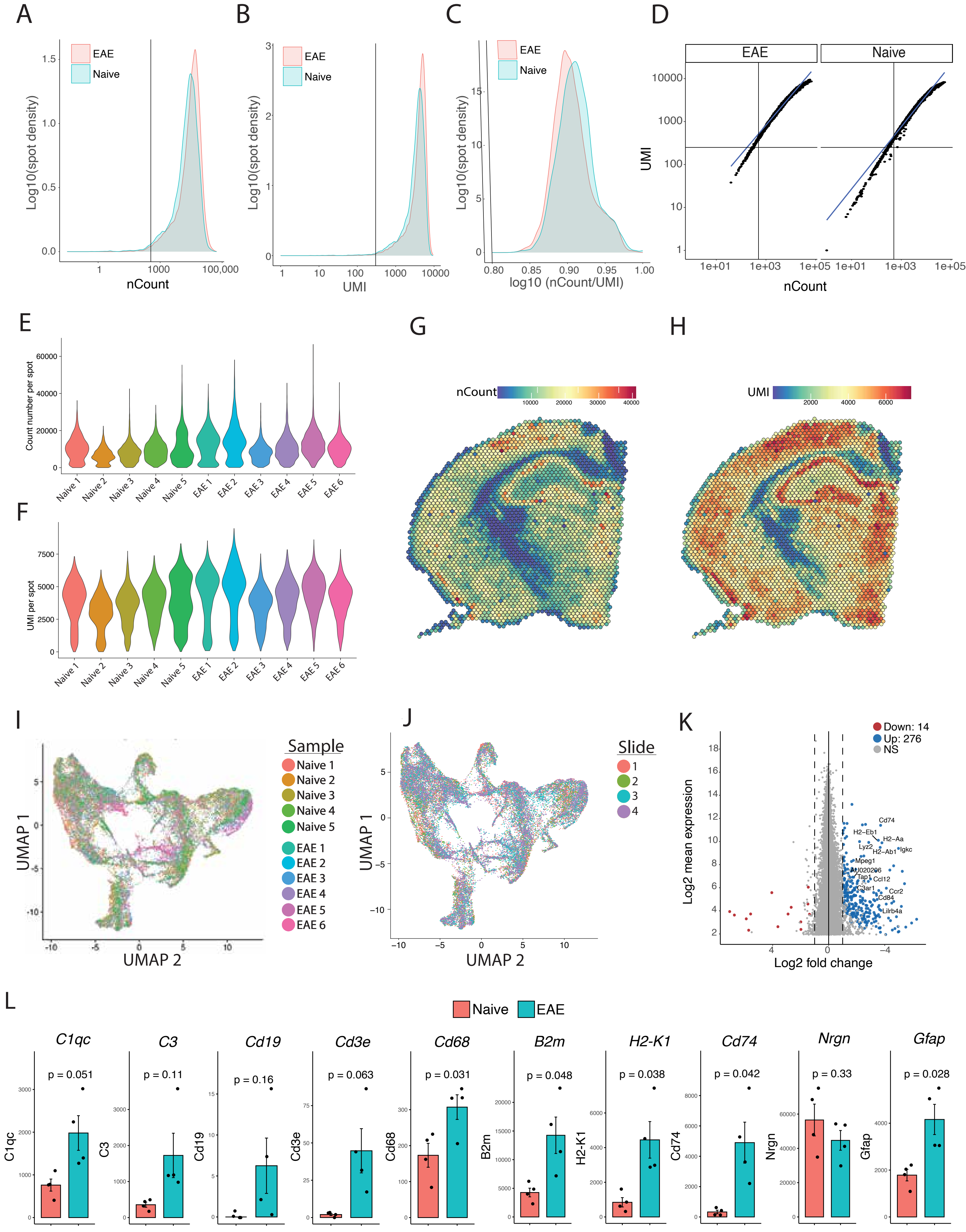

### Supplementary Figure 3

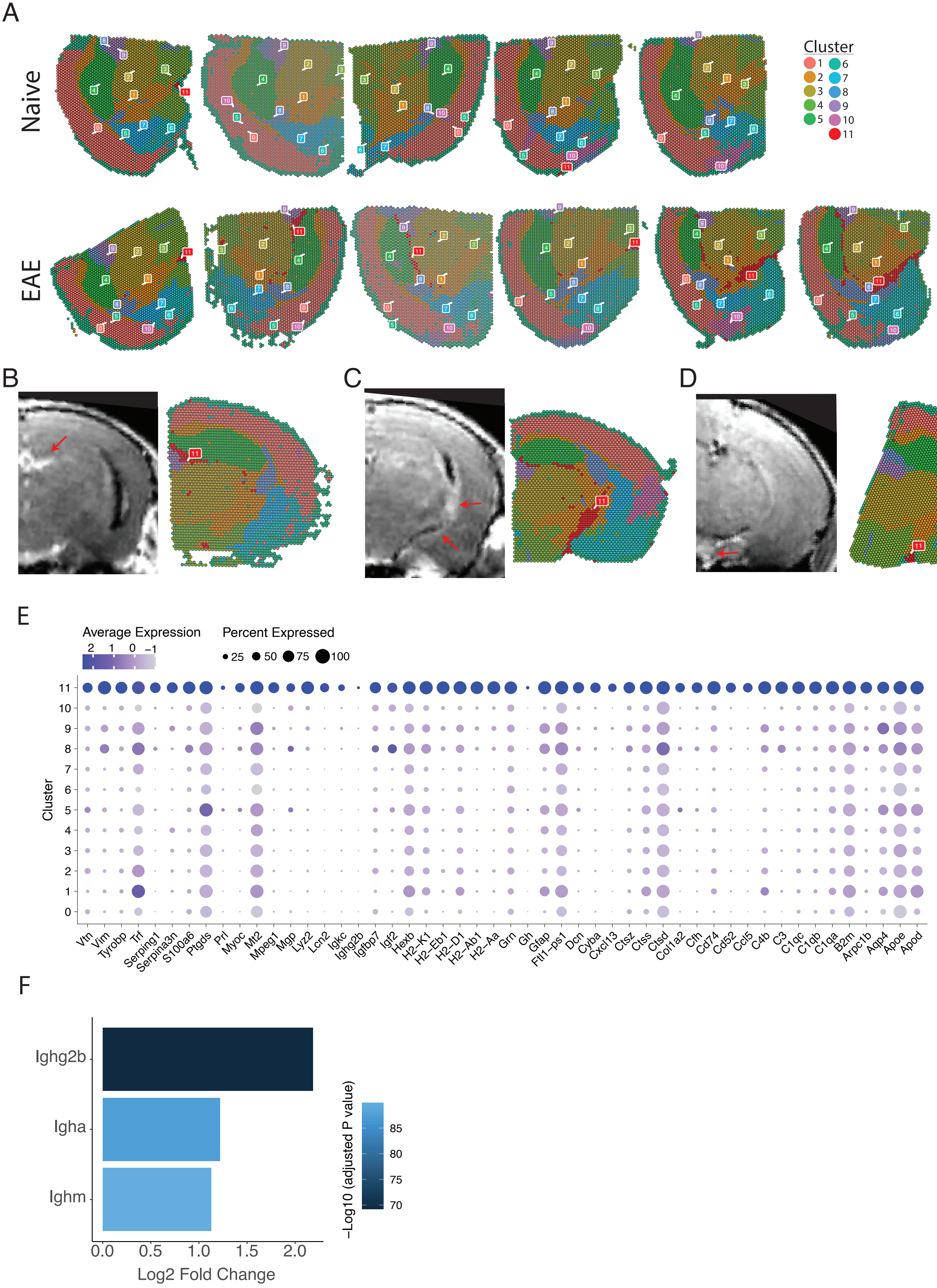

### Supplementary Figure 4

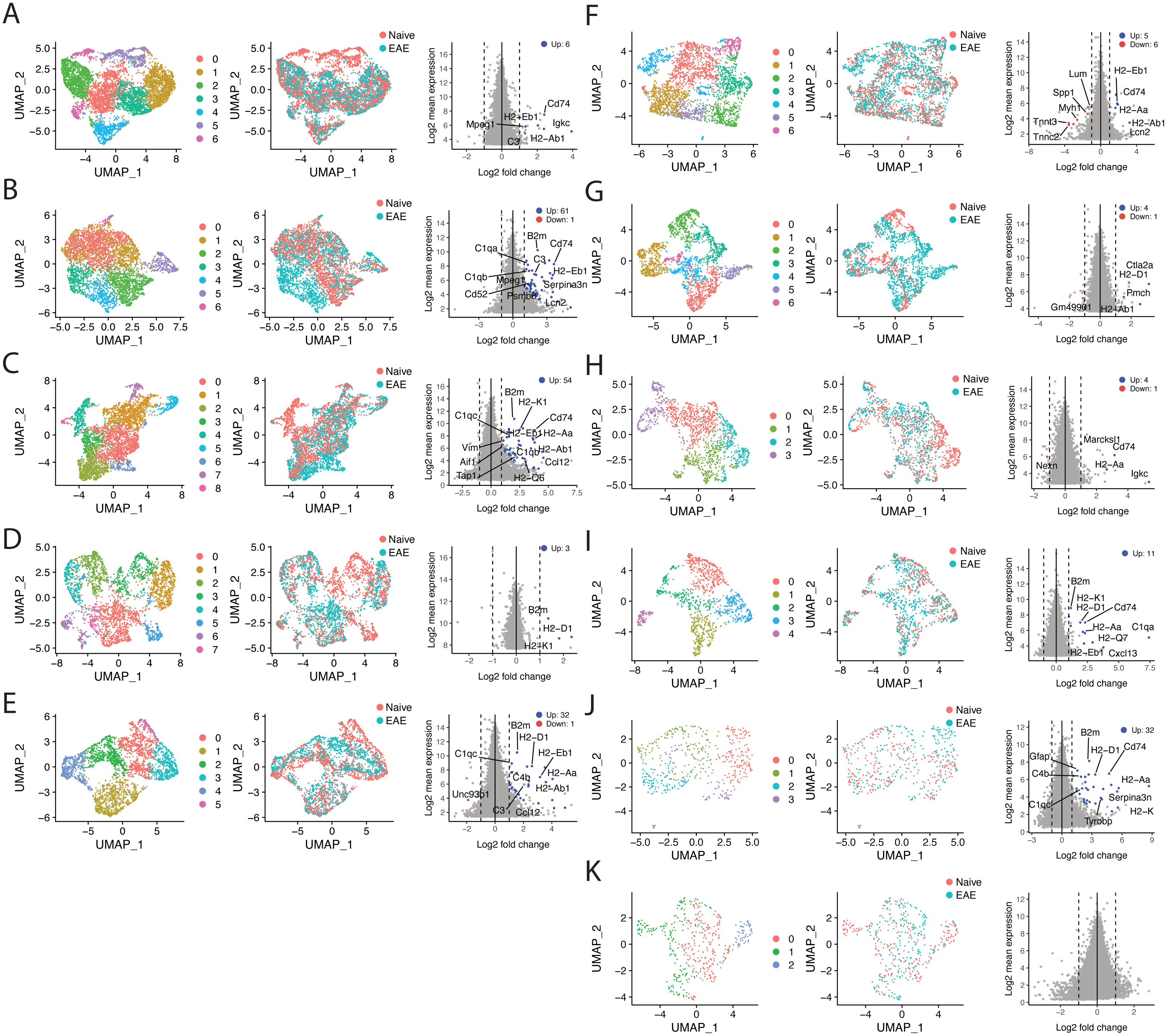

### Supplementary Figure 5

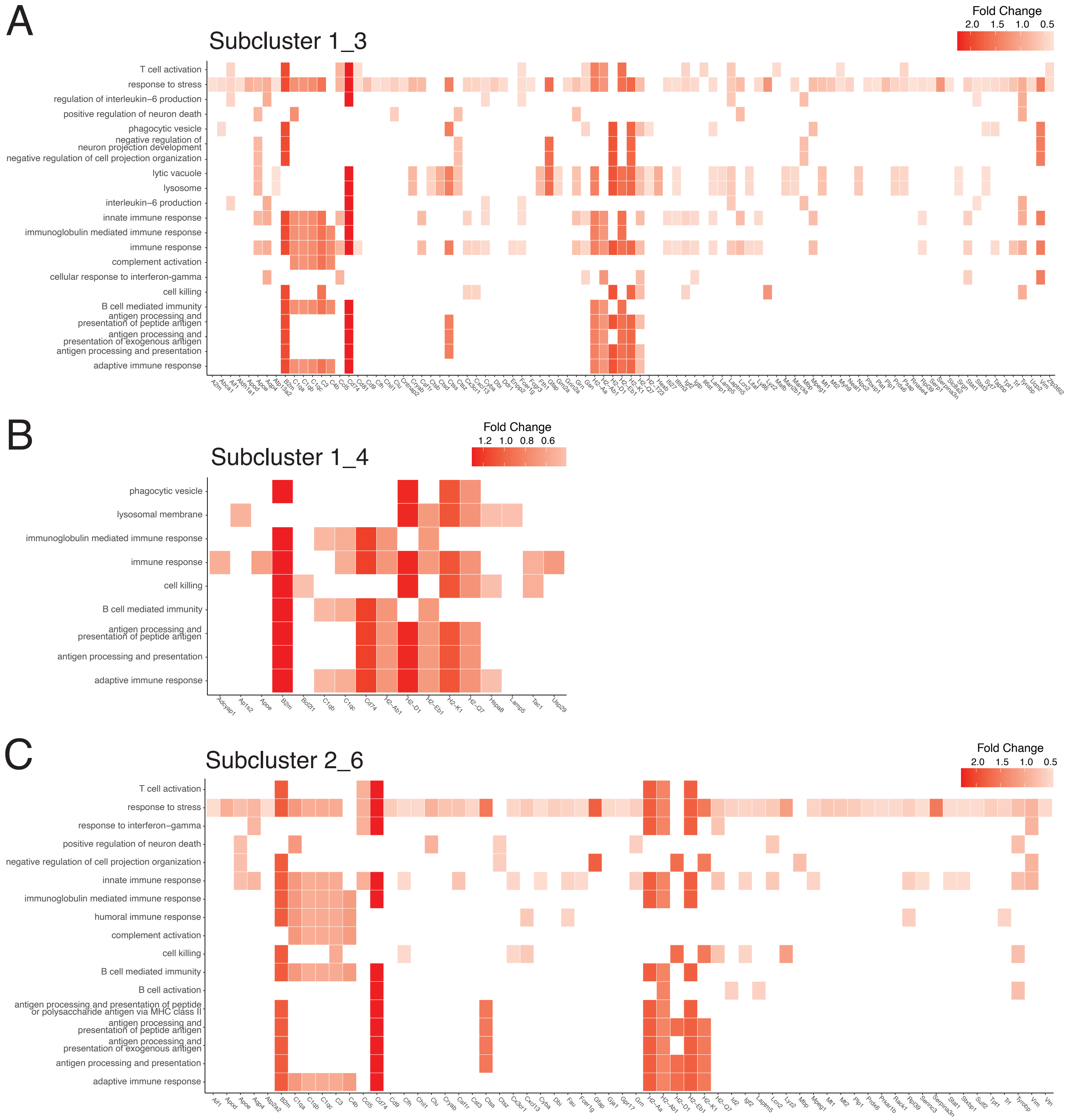

### Supplementary Figure 6

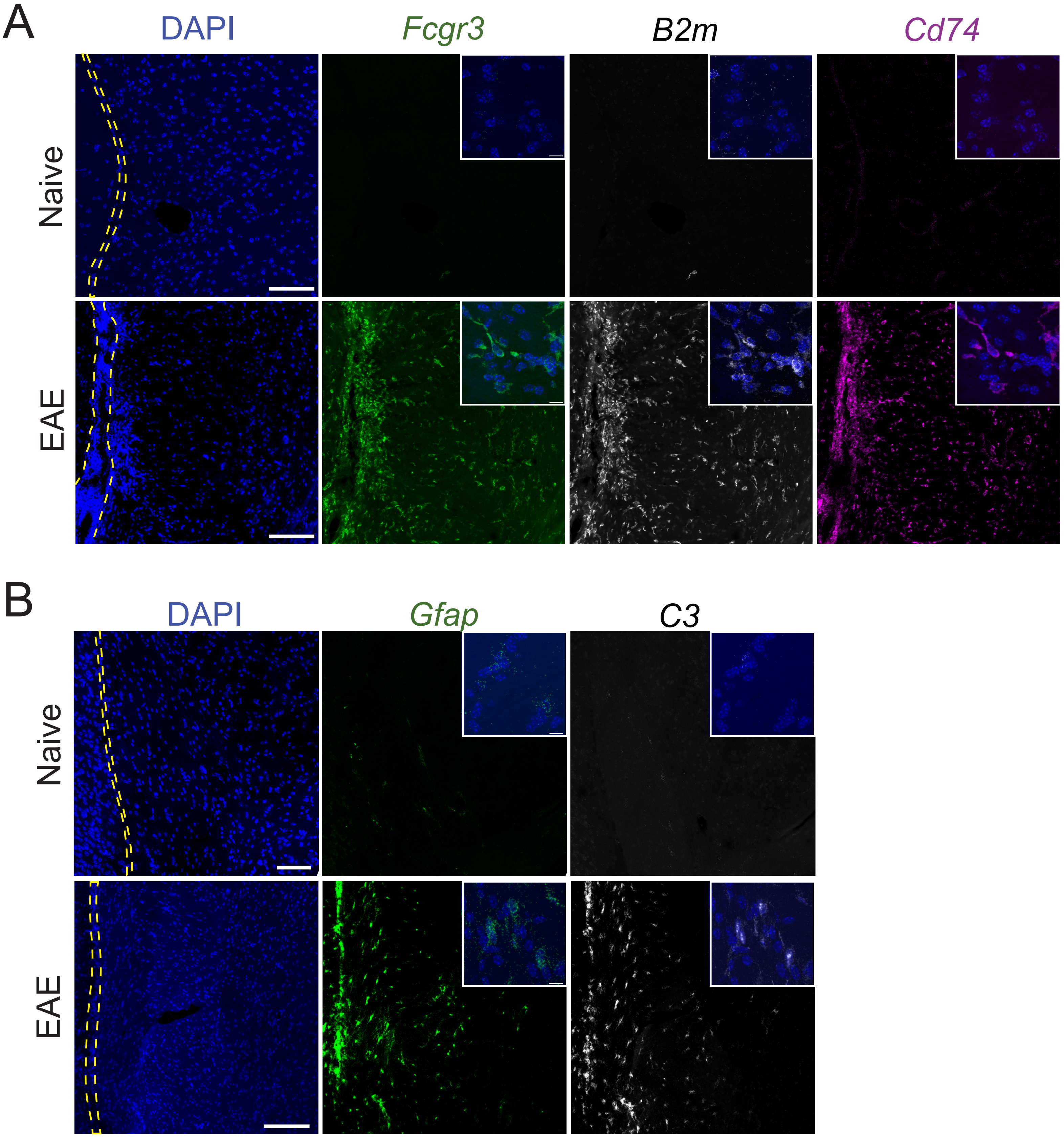
